# TgF344-AD rat model of Alzheimer’s disease: spatial disorientation and asymmetry in hemispheric neurodegeneration

**DOI:** 10.1101/2023.08.14.553199

**Authors:** Boriss Sagalajev, Lina Lennartz, Lukas Vieth, Cecilia Tasya Gunawan, Bernd Neumaier, Alexander Drzezga, Veerle Visser-Vandewalle, Heike Endepols, Thibaut Sesia

**Affiliations:** University of Cologne, Faculty of Medicine and University Hospital Cologne, Department of Stereotactic and Functional Neurosurgery, Cologne, Germany; European Graduate School of Neuroscience (EURON), P.O. Box 5800, 6202 AZ, Maastricht, Netherlands; University of Cologne, Faculty of Medicine and University Hospital Cologne, Institute of Radiochemistry and Experimental Molecular Imaging, Cologne, Germany; Forschungszentrum Jülich GmbH, Institute of Neuroscience and Medicine, Nuclear Chemistry (INM-5), Jülich, Germany; University of Cologne, Faculty of Medicine and University Hospital Cologne, Department of Nuclear Medicine, Cologne, Germany; Forschungszentrum Jülich GmbH, Institute of Neuroscience and Medicine, Molecular Organization of the Brain (INM-2), Jülich, Germany

## Abstract

**BACKGROUND:** The TgF344-AD ratline represents a transgenic animal model of Alzheimer’s disease (AD). We previously reported spatial memory impairment in TgF344-AD rats, yet the underlying mechanism remained unknown. We, therefore, set out to determine if spatial memory impairment in TgF344-AD rats is attributed to spatial disorientation. Also, we aimed to investigate whether TgF344-AD rats exhibit signs of asymmetry in hemispheric neurodegeneration, similar to what is reported in spatially disoriented AD patients. Finally, we sought to examine how spatial disorientation correlates with working memory performance.

**METHODS:** TgF344-AD rats were divided into two groups balanced by sex and genotype. The first group underwent the delayed match-to-sample (DMS) task for the assessment of spatial orientation and working memory, while the second group underwent positron emission tomography (PET) for the assessment of glucose metabolism and microglial activity as in-vivo markers of neurodegeneration. Rats were 13 months old during DMS training and 14-16 months old during DMS testing and PET.

**RESULTS:** In the DMS task, TgF344-AD rats were more likely than their wild-type littermates to display strong preference for one of the two levers, preventing working memory testing. Rats without lever-preference showed similar working memory, regardless of their genotype. PET revealed hemispherically asymmetric clusters of increased microglial activity and altered glucose metabolism in TgF344-AD rats.

**CONCLUSIONS:** TgF344-AD rats display spatial disorientation and hemispherically asymmetrical neurodegeneration, suggesting a potential causal relationship consistent with clinical observations. In rats without spatial disorientation, working memory remains intact.

## Introduction

Alzheimer’s disease (AD) remains incurable despite advances in understanding this condition’s neuropathology and continuous efforts for new therapeutic approaches. One main obstacle to effective therapy is that the brain has already suffered extensive or unredeemable damage when patients are diagnosed. At the diagnostic stage, the AD brain is characterized by the formation of extracellular amyloid plaques (AP) from amyloid-beta (Aβ) peptide fibrils and of intracellular neurofibrillary tangles (NFT) from hyperphosphorylated tau (p-tau) protein [1]. Accumulation of AP and NFT leads to degeneration of neurons and synapses and eventually to the progressive cognitive decline typical of AD [2]. Accordingly, the AP and neuroinflammation correlate with the severity of cognitive impairment in AD patients, while drugs independently targeting both biomarkers fail at yielding significant clinical benefits [3]. AD starts decades before its clinical presentation and diagnosis [4]. Unfortunately, carrying out studies at such an early stage in prospective patients is difficult for practical and ethical reasons. Nevertheless, many challenges to answering pressing questions about the neuropathology of AD can be tackled with suitable transgenic rat models.

The “amyloid cascade hypothesis” states that Aβ-aggregation initiates a chain of pathophysiological reactions that ends in neuronal loss and the development of dementia [5]. The hypothesis is supported by genetic evidence that any mutation in the amyloid precursor protein (APP) or in presenilin 1 or 2 (PS1/PS2) eventually leads to Aβ accumulation and thus to the development of early-onset AD in humans [1]. This line of evidence led to the generation of Aβ-overproducing transgenic mice, which until recently served as the “gold standard” in AD animal modeling, but which did not develop robust tauopathy nor neuronal loss unless additional human transgenes were introduced [6-8]. Rats, on the other hand, are phylogenetically closer to humans with regards to key physiological aspects such as the number of tau isoforms [9, 10]. Therefore, the transgenic rat line TgF344-AD, which was generated from Fischer 344 rats, is one of the most suitable animal models for AD research, as it manifests in an age-dependent manner, similar to patients, the complete repertoire of AD’s pathological hallmarks: cerebral amyloidosis, tauopathy, oligomeric Aβ, gliosis, apoptotic loss of neurons, and behavioral impairment [9]. Specifically, the TgF344-AD rat line expresses two mutant human genes: the “Swedish” APP (*APPsw*) and the Δ exon 9 PS1 (*PS1ΔE9*). Numerous studies have validated the phenotype of TgF344-AD rats [11-14], which, since their first description, have been used to test efficacy of deep brain stimulation [15, 16], Aβ attenuation therapy [17], anti-inflammatory agents [18], and low-dose brain radiation [19]. Converging data suggest that the time resolution of the symptoms’ progression in this transgenic line not only allows for studying AD at both early and advanced stages but also at a prodromal stage [20-23].

In our previous study, we showed that spatial memory is impaired in Tgf344-AD rats, yet the exact mechanism remained unknown [15]. We, therefore, set out to determine if spatial memory deficits in Tgf344-AD rats result from the development of spatial disorientation, which, as in AD patients [2, 24, 25], may result from the asymmetry in hemispheric neurodegeneration. We used positron emission tomography (PET) to detect changes in glucose metabolism and microglial activity as a means of assessing neurodegeneration in vivo [26]. Also, we subjected Tgf344-AD rats to the delayed match-to-sample (DMS) task to assess their ability to differentiate between left and right sides as well as to simultaneously measure their working memory performance.

## Methods

### Ethical approval

All experiments were approved by the local regulatory authority (*Landesamt für Natur, Umwelt und Verbraucherschutz Nordrhein-Westfalen* (LANUV), North Rhine-Westphalia, Germany; permission no. 84-02.04.2015.A490 and 81-02.04.2022.A045) and were carried out in accordance with the EU Directive 2010/63/EU on the protection of animals used for scientific purposes. ARRIVE guidelines on reporting of animal research were followed.

### Animals

A colony of Tgf344-AD rats was bred at the University of Cologne as part of the Materials Transfer Agreement (Rat Resource and Research Center, University of Missouri, Columbia, MO, USA). To avoid any potential effects that carrying the transgenes may have on maternity, we only allowed transgene-positive male rats to mate with wild-type Fischer 344 female rats (Janvier Labs, Le Genest-Saint-Isle, France). Transgenic pups did not show obvious AD signs, so no differences could be made a priori between them and their wild-type littermates. The genotype was, therefore, determined with PCR using ear skin biopsy taken at the time of weaning (Transnetyx Inc., New York, NY, USA).

Rats were divided into two groups balanced by sex and genotype. The first group consisted of 10 transgenic (7♀ + 3♂) and 10 wild-type (5♀ + 5♂) rats, which underwent the DMS task. The second group consisted of 6 transgenic (3♀ + 3♂) and 6 wild-type (3♀ + 3♂) rats, which underwent PET for the measurement of microglial activity, as well as of 5 transgenic (2♀ + 3♂) and 6 wild-type (3♀ + 3♂) rats, which underwent PET for the measurement of glucose metabolism. A slight bias in the ratio of female to male transgenic rats was attributed to the varying number of pups of different sexes per litter. DMS training started when the rats were 13 months old, while DMS testing started when they were 14 months old. PET was performed when the rats were 14-16 months old.

Between the experiments, the rats were kept together in cages littered with low-dust spruce granulate and enriched with cardboard tubes, paper towels, and aspen bricks. While access to food in the PET group was ad libitum, in the DMS group it was restricted to 3-8 h per day. This was done to maintain the rats’ weight at 85%-90% of their ad-libitum value and thus increase their motivation to obtain a sucrose pellet during the DMS task. No restrictions on water intake were imposed for either group.

All experiments were conducted from 9 AM to 6 PM. However, due to different light cycles in different animal facilities, DMS rats (normal 12:12 h) were tested a few hours after dawn, while PET rats (inverse 12:12 h) a few hours after dusk. Rats are crepuscular animals, meaning they are most active during dawn and dusk. Therefore, despite the different light cycles, both the PET and DMS rats were tested when they were least active, making them comparable in this regard. However, to mitigate any potential effects that varying light conditions could have had on our findings, we conducted only within-group analyses.

### DMS task

#### Habituation

Spatial orientation and working memory were evaluated using the DMS task according to a procedure adapted from Dunnett et al. using isolated operant chambers [27]. The chambers had a pellet dispenser and two retractable levers to the left and right of the feeder. Before the DMS task, the rats were handled for five minutes daily for several days to decrease their stress levels. The handling was followed by a habituation period of at least two days when the rats stayed for 30 min inside the operant chambers. Then the training part of the DMS task started.

### Training

The training consisted of five stages. The first four stages familiarized the rat with the mechanism of pressing levers and obtaining a sucrose pellet as a reward (45 mg, unflavored Dustless Precision Pellets; Bio-Serv., Flemington, NJ, USA). Throughout the training stages, an incorrect response (pressing the incorrect lever or failing to press any lever) produced a timeout of 5 s, during which the overhead lights were turned off and no sucrose pellet was delivered. The trials were separated by 10 s. The stages went as follows:

– Stage I: The two levers are extended, and the rat is rewarded for every single lever press.
– Stage II: Only one lever is extended, with the side of the lever being randomized across trials. Once the extended lever is pressed, it retracts and then extends again without delay. The rat is rewarded following the second lever press.
– Stage III: Similar to stage II, except that the active lever changes side every block of three successful trials.
– Stage IV: Similar to stage III, except that the active lever changes side randomly between trials.
– Stage V: The rat must press the lever presented at random on the left or right side to initiate the trial; it is the sample lever for that trial. After the sample lever retracts, both levers extend without delay. The rat is rewarded for pressing (i.e., matching) the sample lever. This stage is similar to the DMS testing part (described below) but without the delay between the sample and match phases.

Every experimental day, stages I-II had 60 trials each, while stages III-IV lasted until the rat had 30 pellets (max one pellet per trial). Finally, stage V ended after 60 minutes or when all 60 trials were completed, whichever came first. Only when the rat achieved a minimum of 75% correct responses in stage V, did it proceed to the testing part of the DMS task.

#### Testing

During the testing part of the DMS task, the delay between sample and match phases was randomly set to 1, 2, 4, 8, or 16 s. Each delay was presented 12 times per experimental day, resulting in a total of 60 trials. The primary measure was the longest delay with a success rate of over 75%, as only then the rat was considered to use directed strategy.

#### Statistics

The statistical analysis was done in Python (version 3.9.13) using SciPy (version 1.9.1) and Pingouin (version 0.5.3) libraries. Figures were made in Python using Matplotlib (version 3.7.0) and Seaborn (version 0.12.2) libraries.

An unpaired two-tailed t-test was used to assess the learning rate and lever preference during DMS training, while a two-way mixed ANOVA with Geisser-Greenhouse (GG) correction was used to assess working memory during DMS testing. In addition, a Fisher’s exact test was conducted to establish the association between lever preference and genotype. Results were considered significant at a p-value below 0.05 only when statistical power (1−β) exceeded 0.8 at a significance level (α) of 0.05.

### PET imaging

#### Image acquisition

[^18^F]FDG, purchased from Life Radiopharma Bonn GmbH (Germany), was used to assess brain glucose metabolism at rest. The rats were briefly anesthetized with isoflurane in O2/air 3:7 (induction 5%, maintenance 2%), and 58-69 MBq [^18^F]FDG in 500 μl was injected intraperitoneally. The rats were then placed in a recovery cage, where they spent the following 50 min awake. Subsequently, they were anesthetized again and fixed with a tooth bar in an animal holder (Minerve, Esternay, France) with a respiratory mask for isoflurane delivery. Rats were warmed by heated airflow to keep body temperature at 37 °C. Their eyes were protected from drying out by applying eye and nose ointment (Bepanthen, Bayer, Germany). The breathing rate was monitored and kept at 40-60 breaths/min by adjusting isoflurane concentration. A static PET scan was conducted using a Focus 220 micro PET scanner (CTI-Siemens, Erlangen, Germany) with a resolution at the center of the field of view of 1.4 mm. The emission scan started 60 min after [^18^F]FDG injection with an acquisition time of 30 min. It was followed by a 10-minute transmission scan using a ^57^Co point source for attenuation correction. This protocol takes advantage of metabolic trapping of [^18^F]FDG [28], which allows awake tracer uptake and subsequent scanning under anesthesia [29].

[^18^F]DPA-714 was synthesized at the Forschungszentrum Jülich GmbH (Germany). The rats were anesthetized with isoflurane in O2/air 3:7 (induction 5%, maintenance 2%), and a catheter for tracer injection was inserted into the lateral tail vein. After fixation in the animal holder, the emission scan started with intravenous injection of 58-67 MBq [^18^F]DPA-714 in 500 μl. Acquisition time was 30 min, followed by a transmission scan as described for [^18^F]FDG. After the scan was finished, the catheter was removed, and the rats woke up in their home cages.

#### Image reconstruction and statistics

After full 3D rebinning (span 3, ring difference 47), summed images were reconstructed using an iterative OSEM3D/MAP procedure [30], resulting in voxel sizes of 0.38 × 0.38 × 0.80 mm. For all further processing of the images including statistics, the software VINCI 4.72 for MacOS X (Max Planck Institute for Metabolism Research, Cologne, Germany) was used. Images were co-registered and intensity-normalized to a reference region. To this end, an elliptical volume of interest (VOI) was placed inside a background region, which was a consistently selected tracer-negative brain area in the midbrain for [^18^F]DPA-714 (4 mm^3^) [31], and the cerebellum for [^18^F]FDG (25 mm^3^) [32]. Each image was divided by the mean value of the reference VOI, resulting in the “standardized uptake value ratio” (SUVR_bg_ for [^18^F]DPA-714 and SUVR_Cer_ for [^18^F]FDG). No further postprocessing (e.g., Gauss filtering or spatial morphing) was done.

For comparison of transgenic vs wild-type rats, a t-test was performed for each tracer at the voxel level, followed by a threshold-free cluster enhancement (TFCE) procedure with subsequent permutation testing [33] to correct for multiple comparisons. Results were considered significant at a p-value below 0.05 only when statistical power (1−β) exceeded 0.8 at a significance level (α) of 0.05.

## Results

### DMS task

Behavioral impairments were assessed using the DMS task divided into two parts: training and testing. During the last stage of the training part, there was no delay between the match and sample phases. Hence, the rats were expected to rely only on their spatial orientation abilities to determine which lever to press, left or right. The criterion for completing the last stage was to achieve a correct response rate of 75%. There was a statistically significant difference in the minimum number of days necessary to complete the last stage between the wild-type and transgenic rats (unpaired two-tailed t-test, t(18) = 2.84, p = 0.01). Specifically, all ten wild-type rats took only five days to complete the last stage, while only half of the ten transgenic rats were able to complete it in the same period (**Fig. 1A, B**). The remaining five transgenic rats could not be trained because they pressed one of the two levers significantly more (**Fig. 1C, D**; unpaired two-tailed t-test, t(18) = -2.45, p < 0.05) and therefore were excluded from the next part of the DMS task, that is, working memory testing. A Fisher’s exact test showed that there was a significant association between lever preference and genotype (p < 0.05).

**Figure 1.**
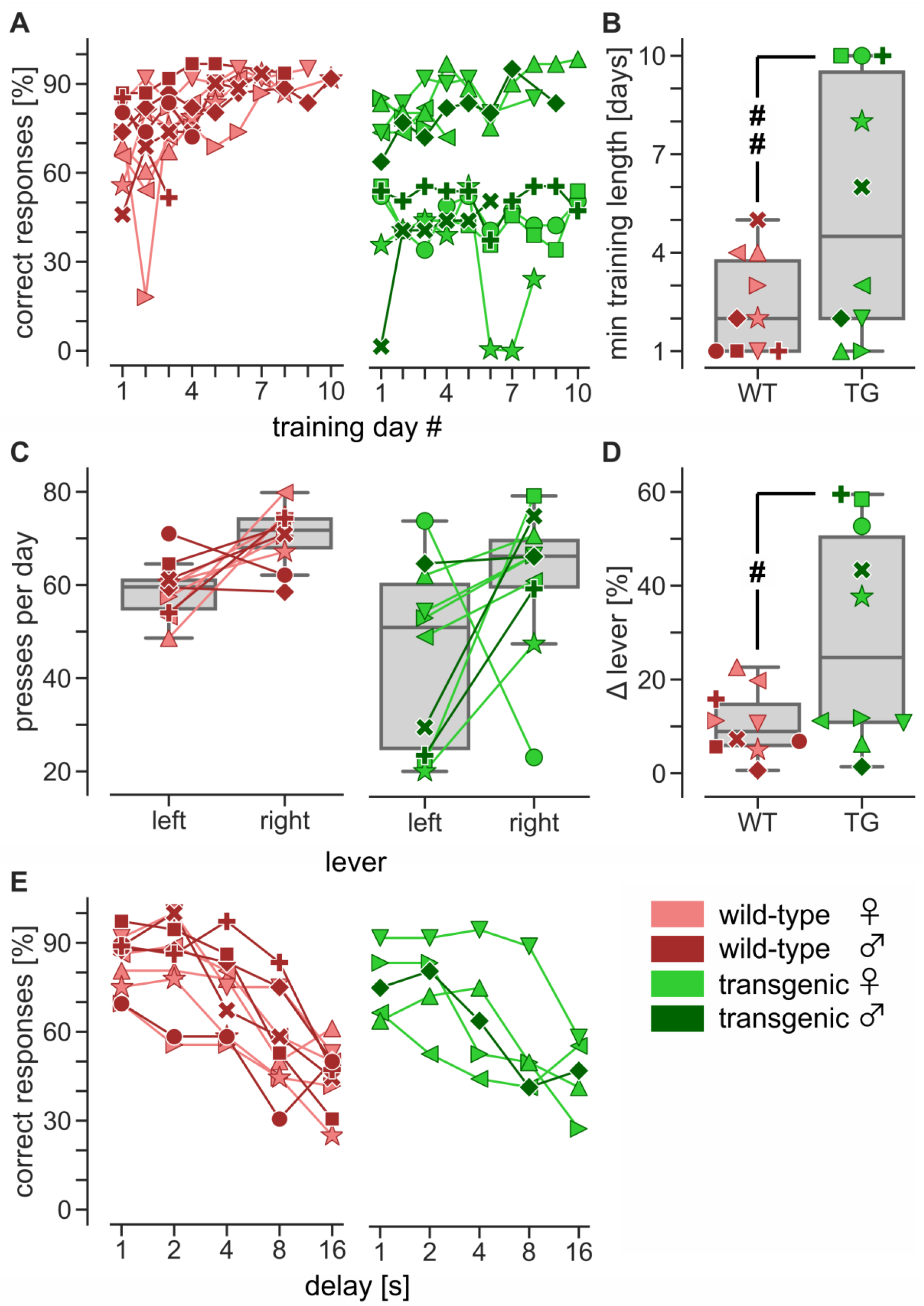
*DMS task*. **A:** Five out of ten transgenic (TG) rats were unable to complete the last stage of the training part, that is, when the delay between the match and sample phases was still absent. **B:** For statistical analysis, their last day of training was used as the minimum length necessary to learn the DMS task (^##^ unpaired two-tailed t-test, t(18) = 2.84, p = 0.01). **C, D:** The difference in the learning rate between transgenic and wild-type (WT) rats was due to a strong preference that the non-learners had for one of the two levers (^#^ unpaired two-tailed t-test, t(18) = -2.45, p < 0.05). **E:** Only rats without a strong preference for either lever proceeded to the testing part where the delay between the match and sample phases was 1-16 seconds. The results of a two-way mixed ANOVA with GG correction showed a decline in performance as the delay increased (F(4, 64) = 33.49, GG corrected p < 0.0001). However, this decline was independent of genotype (genotype: F(1, 16) = 1.68, uncorrected p = 0.21; interaction: F(4, 64) = 1.06, uncorrected p = 0.38). Different symbols are used to show individual rats.

During working memory testing, a delay between the match and sample phases was introduced. In each trial, the delay was randomly set to either 1, 2, 4, 8, or 16 s. We averaged the number of correct responses for each delay every three consecutive days to smoothen the curve depicting rats’ daily performance. We then determined the peak performance of each rat and compared it between the groups (**Fig. 1E**). We compared peak performance rather than final performance to exclude data collected after the rat lost interest in the task. A two-way mixed ANOVA with GG correction was performed to assess the effect of genotype (between-subjects factor) and delay (within-subjects factor) on performance. It showed that the interaction between the effects of genotype and delay was statistically insignificant (F(4, 64) = 1.06, uncorrected p = 0.38). Likewise, the main effect of genotype on performance was statistically insignificant (F(1, 16) = 1.68, uncorrected p = 0.21). In contrast, the main effect of delay was significant (F(4, 64) = 33.49, GG corrected p < 0.0001). Our results, therefore, show that performance declined as delay increased, yet the decline in performance was independent of genotype.

### PET imaging

PET imaging with the TSPO ligand [^18^F]DPA-714 showed confined clusters of increased microglial activity in transgenic rats compared to wild-type rats, with no significant global changes observed (**Fig. 2A**). Bilateral clusters were found in the dorsal subiculum and temporal association cortex, while unilateral clusters were found in the retrosplenial cortex (left), reticular formation (left), ventral thalamus (right), lateral posterior thalamus (left), and hippocampus (left), with the latter having the highest SUVR_bg_ (1.21 ± 0.06 in transgenic rats vs 1.03 ± 0.06 in wild-type rats).

**Figure 2.**
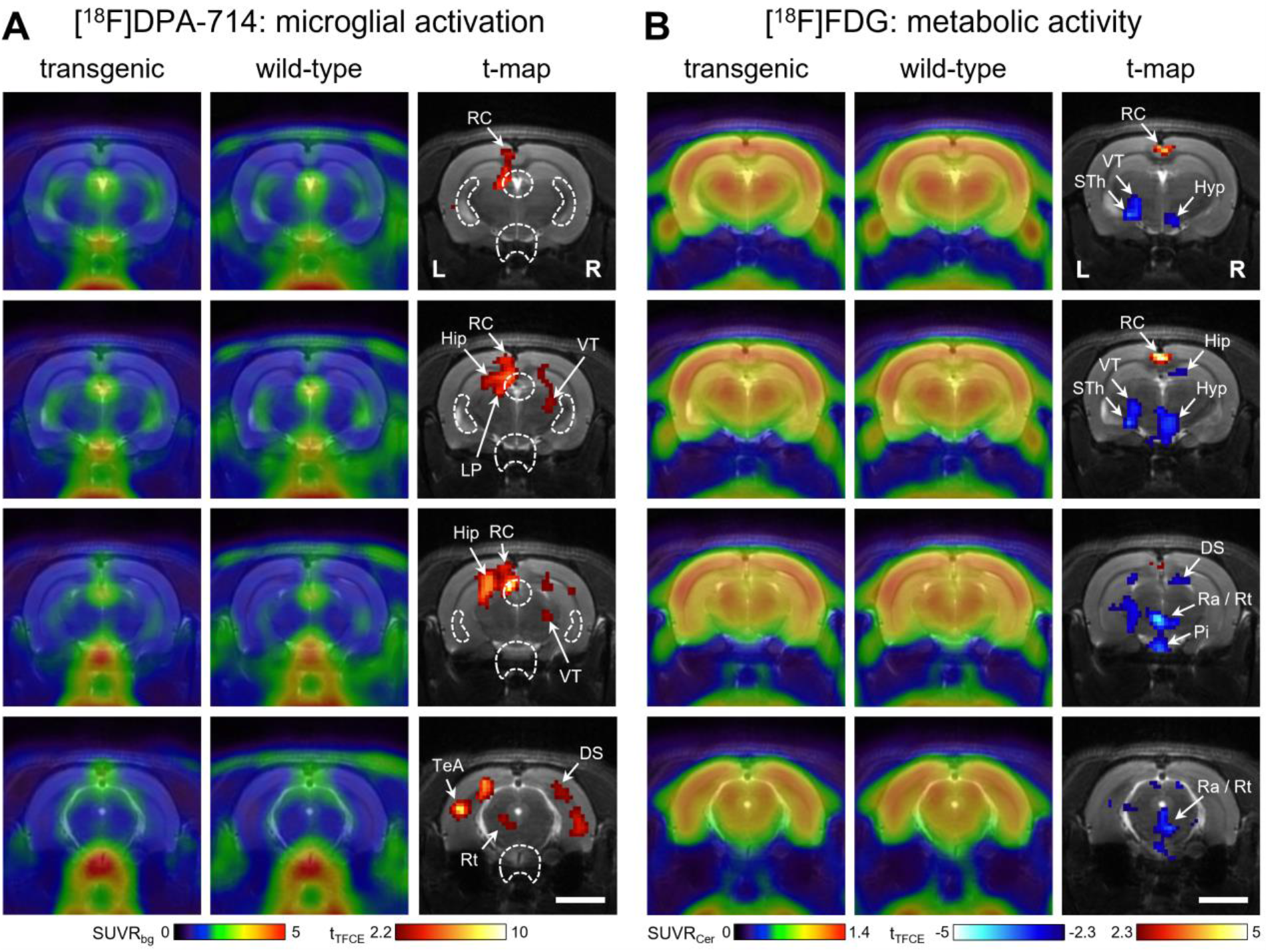
*PET imaging*. Shown are averaged PET images of (**A**) microglial activation measured with [^18^F]DPA-714 (6 transgenic and 6 wild-type rats) and (**B**) glucose metabolism measured with [^18^F]FDG (5 transgenic and 6 wild-type rats). Respective t-maps show significant differences (corrected for multiple testing) between groups. Red voxels: higher tracer uptake in transgenic rats. Blue voxels: higher tracer uptake in wild-type rats. White dashed lines in **A** indicate regions obscured by spillover from the ventricular system and pituitary gland, where natural TSPO expression leads to high tracer binding. DS – dorsal subiculum, Hip – hippocampus, Hyp – hypothalamus, LP – lateral posterior thalamus, Pi – pituitary gland, Ra – raphe, RC – retrosplenial cortex, Rt – reticular formation, STh – subthalamic nucleus, TeA – temporal association cortex, VT – ventral thalamus. Scale bar: 5 mm.

Similarly, PET imaging with [^18^F]FDG showed confined clusters of altered glucose metabolism in transgenic rats compared to wild-type rats, with no significant global changes observed (**Fig. 2B**). Hypometabolic clusters were found in the dorsal subiculum (right), hippocampus (right), hypothalamus (right), ventral thalamus (left), subthalamic nucleus (left), and pituitary gland (bilateral). The most pronounced hypometabolic cluster was found in the reticular formation (bilateral), with an SUVR_Cer_ of 0.88 ± 0.03 in transgenic rats vs 0.99 ± 0.05 in wild-type rats. The only hypermetabolic cluster was found in the retrosplenial cortex (bilateral), with an SUVR_Cer_ of 1.21 ± 0.03 in transgenic rats vs 1.13 ± 0.03 in wild-type rats.

## Discussion

Our behavioral data demonstrate that half of Tgf344-AD rats developed a clear preference for one of the two levers (left vs right) during the training phase of the DMS task (delay = 0 s), indicating the presence of spatial disorientation. This is consistent with the results of our previous study showing that Tgf344-AD rats exhibit spatial memory deficits in a modified Barnes maze, which can be attenuated by deep brain stimulation [15]. Interestingly, our present study also shows that Tgf344-AD rats without spatial disorientation perform equally well as their wild-type littermates in the testing phase of the DMS task (delay = 1-16 s), suggesting that their working memory is intact. This finding is consistent with the results of Muñoz-Moreno et al., who used the nonmatch version of the same task and showed that TgF344-AD rats exhibit a similar percentage of correct responses as their wild-type counterparts across all delays [34]. In a novel object recognition (NOR) test, TgF344-AD rats are reported to show no side bias [20], which somewhat contradicts our results of spatial disorientation in the DMS task. However, the reason for this difference is likely because in the NOR test, rats can face objects from any direction, while in the DMS task, rats can face levers only from one direction. This ensured that, in our DMS task, the left lever always remained on the left, while the right lever on the right.

In a transgenic mouse model of AD, whole-brain glucose hypermetabolism was shown to precede glucose hypometabolism [35, 36], which in turn is shown to positively corelate with microglial activity in early-onset AD patients [26]. Activation of microglia and astrocytes around AP is classically regarded to mark the onset of neuroinflammation in AD [37]. Our PET data on resting glucose metabolism and microglial activity suggest a mosaic of focal changes, which is in line with magnetic resonance imaging studies showing disruption of structural and functional networks in TgF344-AD rats as early as 5 months of age, confirming the notion of AD as a disconnection syndrome [34, 38].

Our PET results show an increase in microglial activity in the hippocampus, dorsal subiculum, thalamic nuclei, as well as in the temporal association and retrosplenial cortices of Tgf344-AD rats. This increase is consistent with another PET study that used the same TSPO ligand (i.e., [^18^F]DPA-714), showing no change in TgF344-AD rats at 6 months of age, yet an increase in neuroinflammation at 12 months in the hippocampus and at 18 months in the thalamus and frontal cortex [20]. In the hippocampus of TgF344-AD rats, TSPO overexpression was previously reported at 12 months of age in astrocytes, and at 24 months of age in microglia, supporting the notion of TSPO as a neuroinflammatory marker [39]. Our PET findings are also consistent with existing magnetic resonance spectroscopy (MRS) data on increased neuronal dysfunction in the hippocampus of TgF344-AD rats. However, while the MRS data are based on a decrease in N-acetyl-aspartate levels occurring at 18 months of age [20], our study found neuronal dysfunction in the form of impaired glucose metabolism and increased microglial activity in TgF344-AD rats that were 14-16 months of age.

Of relevance in our study are hypometabolic changes in the hypothalamus and pituitary gland of Tgf344-AD rats, as they support the recent hypothesis that dysregulation of the hypothalamic-pituitary-adrenal axis may help in the diagnosis of prodromal AD [40]. Also important in our study is the coincidence of spatial disorientation with increased microglial activity and altered glucose metabolism in the thalamus and retrosplenial cortex since dysfunction of these two interdependent structures represents one of the earliest signs of prodromal AD in patients [41]. Similarly, in a mouse model of amyloid pathology, dysfunction of the thalamus and retrosplenial cortex precedes overt AP formation [42].

Asymmetric neuroinflammation has previously been reported in Aβ mouse models (expressing mutant human APP alone or in combination with PS1/PS2) and was accompanied by fibrillary amyloidosis in the affected areas [43]. The asymmetry in our PET data is in line with clinical observations that brain atrophy in AD patients is bilateral but not always symmetrical [2, 24, 25, 44]. Therefore, some AD patients may develop atrophy predominantly in one of the hemispheres. Often such patients have contralateral spatial neglect, which is not compensated by the healthier hemisphere since it is also partially affected by neurodegeneration [24]. This is consistent with our PET findings, showing that unilateral microglial activation in TgF344-AD rats is accompanied by contralateral glucose hypometabolism in structures such as the ventral thalamus and hippocampus.

In conclusion, TgF344-AD rats display spatial disorientation and hemispherically asymmetrical neurodegeneration, with past clinical research suggesting a causal relationship between the two [2, 24, 25, 44]. When, however, TgF344-AD rats do not develop spatial disorientation, their working memory remains intact.

## Limitations

Although wild-type rats may exhibit functional hemispheric asymmetry [45] that can progress with age [46], our data show not that hemispheric asymmetry is absent in wild-type rats, but only that it is more pronounced in transgenic rats. Also, one has to exercise caution when establishing an association between rats used for the DMS task and PET imaging, as the former, but not the latter, were subjected to food restriction known to enhance cognitive abilities [47]. Therefore, food restriction as a prerequisite of the DMS task may have actually decreased its sensitivity to detect behavioral changes. Additionally, the resting glucose metabolism in PET doesn’t necessarily reflect brain activity in the DMS task. Moreover, while the rats were awake during the DMS task and PET measurement of resting glucose metabolism, they were anesthetized during PET measurement of microglial activity. Studies indicate that isoflurane can provide neuroprotection by alleviating microglial activation [48, 49]. Therefore, microglial activation in TgF344-AD rats may have been stronger if they were awake.

## Acknowledgments

The authors are thankful to Julia Mauz for her support in the data collection process.

## Data Availability

Data supporting the findings of this study are available on request from the corresponding author.

